# MOLI: Multi-Omics Late Integration with deep neural networks for drug response prediction

**DOI:** 10.1101/531327

**Authors:** Hossein Sharifi-Noghabi, Olga Zolotareva, Colin C. Collins, Martin Ester

## Abstract

**Motivation:** Historically, gene expression has been shown to be the most informative data for drug response prediction. Recent evidence suggests that integrating additional omics can improve the prediction accuracy which raises the question of how to integrate the additional omics. Regardless of the integration strategy, clinical utility and translatability are crucial. Thus, we reasoned a multi-omics approach combined with clinical datasets would improve drug response prediction and clinical relevance.

**Results:** We propose MOLI, a **M**ulti-**O**mics **L**ate **I**ntegration method based on deep neural networks. MOLI takes somatic mutation, copy number aberration, and gene expression data as input, and integrates them for drug response prediction. MOLI uses type-specific encoding subnetworks to learn features for each omics type, concatenates them into one representation and optimizes this representation via a combined cost function consisting of a triplet loss and a binary cross-entropy loss. The former makes the representations of responder samples more similar to each and different from the non-responders, and the latter makes this representation predictive of the response values. We validate MOLI on in vitro and in vivo datasets for five chemotherapy agents and two targeted therapeutics. Compared to state-of-the-art single-omics and early integration multi-omics methods, MOLI achieves higher prediction accuracy in external validations. Moreover, a significant improvement in MOLI’s performance is observed for targeted drugs when training on a pan-drug input, i.e. using all the drugs with the same target compared to training only on drug-specific inputs. MOLI’s high predictive power suggests it may have utility in precision oncology.

**Availability of the implemented codes:** https://github.com/hosseinshn/MOLI

## 1 Introduction

Precision oncology is the use of genomic data to tailor therapy for an individual cancer patient. However, response to a cancer treatment—chemotherapy or targeted drugs—is a complex phenotype and often depends on multiple factors especially the genomic profile of the patient (Lee *et al.*, 2018). Presently, only 11% of patients treated with precision oncology can be placed in clinical trials and only 5% of patients benefit from precision oncology (Zehir *et al.*, 2017; Cheng *et al.*, 2018; Marquart *et al.*, 2018). Although there are many reasons underlying this modest success rate, improved drug response prediction will significantly increase the number of patients who benefit from targeted therapy (Marquart *et al.*, 2018) or chemotherapy, and avoid adverse side effects who will not (Gavan *et al.*, 2018; Mishra and Verma, 2010). Various *in vitro* studies of cancer cell lines and patient-derived xenograft mice models (PDX) (Gao *et al.*, 2015) have created datasets such as Genomics of Drug Sensitivity in Cancer (GDSC) (Iorio *et al.*, 2016) and Cancer Cell Line Encyclopedia (CCLE) (Barretina *et al.*, 2012). These datasets provide researchers with multi-omics profiles—consisting of genomic (somatic mutation and Copy Number Aberration or CNA), transcriptomic, proteomic, and methylomic data—together with the response to a large number of targeted and chemotherapy drugs. This is different from patient datasets, which record the response only to one or a few drugs that have been administered to a patient. These *in vitro* datasets enable researchers to investigate the drug response mechanism at a large scale, in particular for many drugs, and all the way from various types of pre-clinical models to patients (Iorio *et al.*, 2016; Barretina *et al.*, 2012). Complementing *in vitro* studies, *in silico* studies have aimed at building computational methods that analyze the cumulative effects of single- or multi-omics data to accurately predict drug response (Geeleher *et al.*, 2014; Ding *et al.*, 2018). These studies usually measure the drug response as the drug concentration that reduces viability by 50% (IC50).

A critical challenge in drug response research is the clinical utility, i.e. whether the outcome of the study is translatable to actual patients (Geeleher *et al.*, 2014, 2017). Ideally to achieve translatability, a computational method should be trained on *in vivo* data, however, available *in vivo* datasets such as The Cancer Genome Atlas (TCGA) datasets (Weinstein *et al.*, 2013) do not have enough patient records with drug response information and in particular, unlike cell line datasets such as GDSC, they do not report responses to multiple drugs. For *in silico* drug response prediction, translatability in the simplest case means that a model with good performance (e.g., high prediction accuracy) on *in vitro* data—trained on more samples compared to *in vivo* data—should also have good performance on *in vivo* data.

The majority of studies suggest that gene expression data are the most effective data type for drug response prediction (Iorio *et al.*, 2016; Geeleher *et al.*, 2014; Ding *et al.*, 2016; Graim *et al.*, 2018). Geeleher et al. (Geeleher *et al.*, 2014) showed that a ridge regression model trained on GDCS gene expression data is translatable to Docetaxel, Cisplatin, Erlotinib, and Bortezomib clinical trial data. They also showed that, for Docetaxel, including non-breast cancer cell lines in model training increases the predictive power of the final model compared to the model only trained on breast cancer cell lines. This ridge regression-based pipeline on gene expression also imputed the drug response for The Cancer TCGA (Weinstein *et al.*, 2013; Geeleher *et al.*, 2017). Despite the predictive power of gene expression, adding other omics data types can increase the predictive power especially in pan-cancer models (Iorio *et al.*, 2016).

Multi-omics data provide a machine learning model with different views of the same sample and promise better characterization of biological processes (Argelaguet *et al.*, 2018; Wang *et al.*, 2014). Multi-omics data have been exploited for different problems such as driver gene identification (Dimitrakopoulos *et al.*, 2018; Mo *et al.*, 2013; Shrestha *et al.*, 2017; Singh *et al.*, 2019), patient stratification (Khakabimamaghani and Ester, 2016), survival prediction (Chaudhary *et al.*, 2018), subgroup discovery (Liang *et al.*, 2015), and drug response prediction (Ding *et al.*, 2018). For the drug response prediction, Ding et al. (Ding *et al.*, 2018) proposed a method that concatenates mutation, CNA, and gene expression data and applies autoencoders to learn features for the concatenated multi-omics cell line data. The learned features were used as the input of an elastic net classifier which predicts the binarized IC50 values. We note that the classifier was validated only on CCLE cell lines without studying its translatability to patients or PDX models.

A critical challenge in multi-omics data analysis is how to integrate different data types. There are two major approaches to multi-omics integration: early integration and late integration (Rappoport and Shamir, 2018). In early integration, all omics data types available for a sample are first concatenated, and then an integrated representation of the sample is created by applying some feature learning method, such as autoencoders (Goodfellow *et al.*, 2016), to that representation. Early integration has three disadvantages: first, it disregards the unique distribution of each omics data type. Second, it requires proper normalization to avoid giving more weight to the omics data type with more dimensions. Third, it further increases the dimensionality of the input data which often is already a challenge for single-omics input data (Rappoport and Shamir, 2018). In late integration, features are learned separately for each omics data type, and these features are then integrated into one unified representation to be used as the input for a classifier or a regressor. The advantage of this approach is that it works with the unique distribution of each omics data type, it can employ single-omics normalization for each data type, and it does not increase the dimensionality of the input space.

In this paper, we explore the problem of drug response prediction and propose MOLI, a **M**ulti-**O**mics **L**ate **I**ntegration method based on deep neural networks. MOLI takes somatic mutation, CNA, and gene expression data as input, and predicts the response to a given drug as the output. MOLI learns features for each omics data type by type-specific encoding subnetworks and concatenates the learned features into one representation of the multi-omics profiles. To the best of our knowledge, MOLI is the first end-to-end late integration method with deep neural networks that optimizes this representation via a combined cost function consisting of a triplet loss function (Schroff *et al.*, 2015) and a binary cross-entropy loss function. The former makes the representations of responder cell lines more similar to each and different from the representations of non-responder cell lines and the latter makes this representation predictive of the IC50 values. As another contribution, MOLI employs transfer learning to increase the size of the training dataset. It trains a drug response model on pan-drug inputs (using all the drugs with the same target) instead of drug-specific inputs. Figure 1 illustrates the workflow of MOLI.

**Fig. 1.**
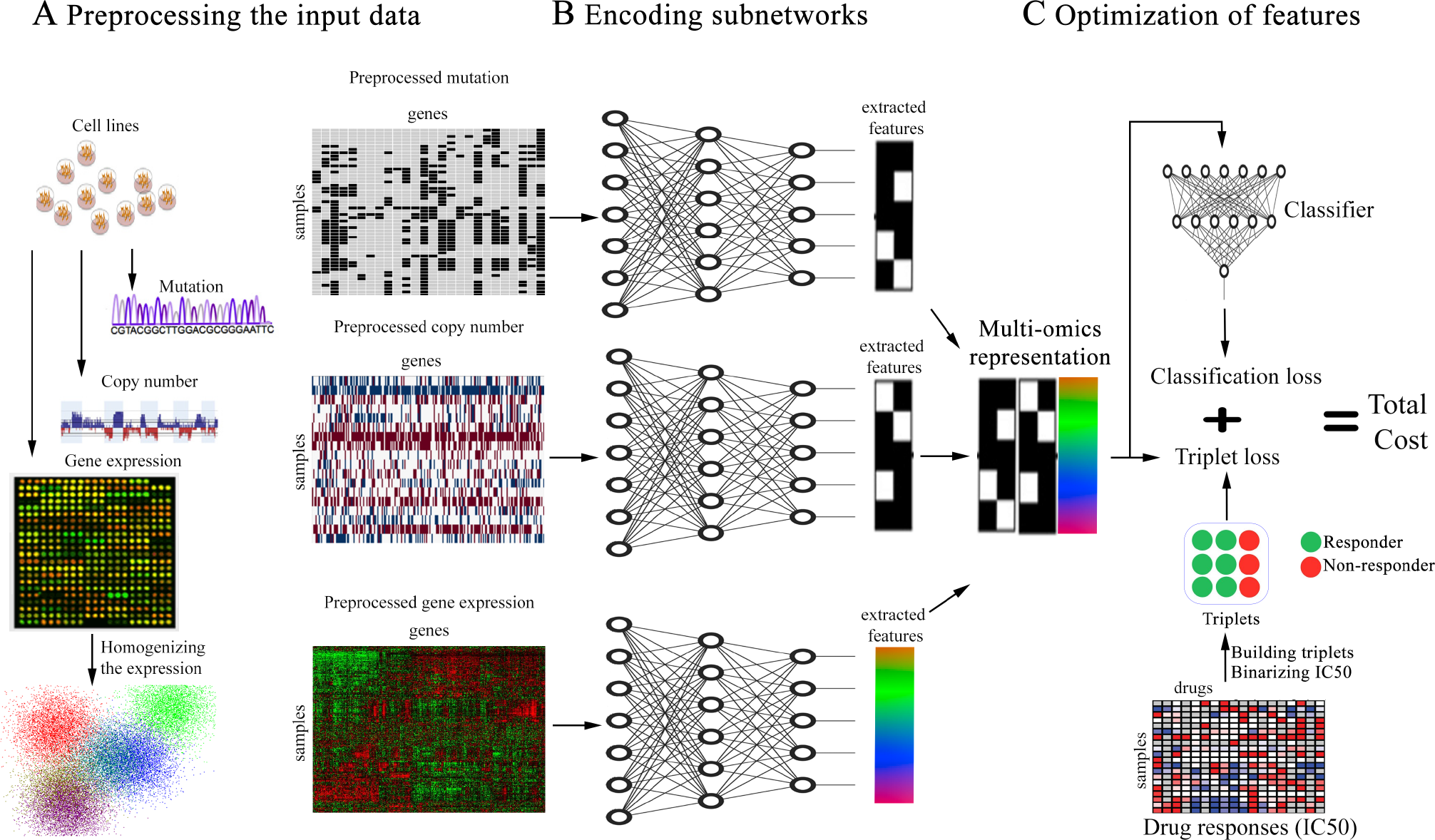
Schematic overview of MOLI (A) preprocessing mutation, CNA, and gene expression data. (B) Each encoding subnetwork learns features for its omics data type and the learned features are concatenated into one representation. (C) MOLI cost function consists of a triplet loss and a classification loss, obtained from the classifier subnetwork that uses the multi-omics representation to predict drug response.

We validated MOLI on *in vitro* (PDX) and *in vivo* (TCGA patients) datasets for five chemotherapy agents and two targeted therapeutics. Our comparison with the state-of-the-art single-omics and early integration multi-omics methods showed that MOLI can achieve significantly better performance in terms of Area Under the receiver operating characteristic Curve (AUC) on PDX and patient data. Moreover, we observed substantial improvement in MOLI’s performance for targeted drugs when training on a pan-drug dataset compared to training on drug-specific datasets. We conclude that MOLI models trained on *in vitro* data translate well to *in vivo* data and may have utility for precision oncology. Finally, we showed that the responses predicted by MOLI—while trained on the pan-drug input for the Epidermal Growth Factor Receptor (EGFR) inhibitors—for breast, lung, kidney, and prostate cancers from TCGA patients (without recorded drug response) had statistically significant associations with some of the genes in the EGFR pathway. This shows that MOLI captures biological aspects of the response.

## 2 Methods

### 2.1 MOLI: Multi-Omics Late Integration

MOLI is a deep neural network that predicts the drug response for a given sample, represented by its multi-omics profile, and for a given drug. MOLI assumes that values for the same genes are provided for each omics data type. MOLI’s network consists of the following subnetworks. It has multiple feed forward encoding subnetworks, one for each input omics data type. Each encoding subnetwork receives its corresponding omics data and encodes it into a learned feature space. The learned features from the encoding subnetworks are integrated into one representation by concatenation. The concatenated representation serves as input for a classification subnetwork, which predicts the drug response. The entire network is trained in an end-to-end fashion using a cost function combining a classification loss and a triplet loss. Figure 1 shows MOLI’s components during training and model development, while Figure 2-A shows the application of MOLI for external validation.

**Fig. 2.**
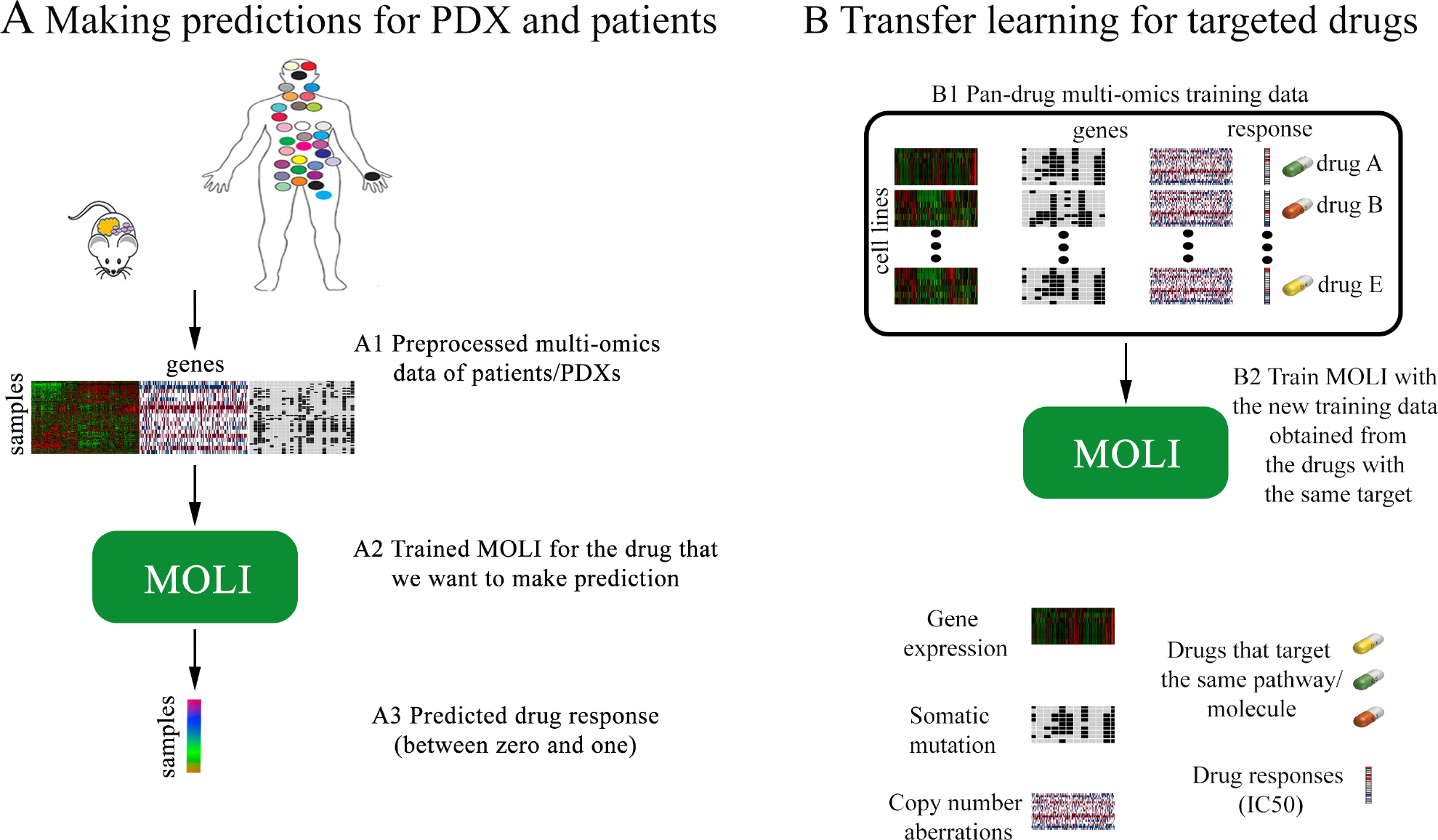
(A) Using MOLI to make predictions for PDX/patient inputs during external validation. (B) Combining targeted drugs that target the same pathway or molecule to make a pan-drug training dataset for MOLI

#### 2.1.1 Learning features by encoding subnetworks

To learn features for each omics data type in the input, we design separate encoding feed forward subnetworks to map the input space to the feature space. In this paper we focus on mutation, CNA, and gene expression data. *X*_*M*_, *X*_*E*_, and *X*_*C*_ denote mutation, CNA, and gene expression data, respectively, each of which are of dimensionality *N* × *D*, where *N* is the number of samples and *D* is the number of genes. We note that the proposed approach can be extended for any number of omics data types. Each encoding subnetwork has a fully connected layer with Relu activation functions. In addition, each subnetwork employs dropout to regularize the model and batch normalization to enhance the training process. The input of each encoding subnetwork is one omics data type and the output is the learned features for that omics (Figure 1-B). We denote these subnetworks as *f*_*M*_ (*X*_*M*_), *f*_*C*_ (*X*_*C*_), and *f*_*E*_ (*X*_*E*_), respectively.

#### 2.1.2 Integrating learned features by late integration

In the integration step, we utilize a late integration approach and concatenate the learned features of the different single-omics data types to obtain one multi-omics representation. For example, if the outputs of three encoding subnetworks are three *M* × *N* feature matrices, after concatenation, the output will be one *M* × 3*N* representation matrix. The integrated representation is further smoothed through a *l*2 normalization layer. We denote MOLI’s integration, receiving multi-omics data as input and returning the integrated representation, as follows:

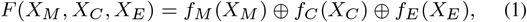

where, ⊕ denotes the concatenation operator.

#### 2.1.3 Optimizing the learned features by the combined cost function

The learned features will be used by a classifier that predicts the drug response. Therefore, the last subnetwork of MOLI is a classification layer with the Sigmoid activation function, using dropout and weight decay for regularization (Figure 1-C). We denote this classifier as *g(.)*. Since the MOLI network will be used for classification, i.e. drug response prediction, the cost function used for training must include a term that measures the difference between the predicted drug response and the ground truth drug response. We choose the binary cross-entropy classification loss, one of the most common classification losses, defined as follows:

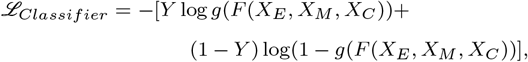
where, *Y*_*N*×1_ denotes the binarized IC50 which is used as measure for the drug response.

We add a triplet loss to the cost function to impose a further constraint that is necessary for accurate classification. This constraint forces responders to be more similar to each other than to non-responders. The triplet loss function was introduced in FaceNet (Schroff *et al.*, 2015) for optimizing the mapping from a space of face images to a Euclidean space where the difference between learned features is correlated with the similarity among faces. The idea is that for the image of a given person’s face, the distance between that image’s learned features and the features of another image of the same person should be smaller than the distance between that image’s learned features and the learned features of the image of some other person. In our context, we employ the triplet loss function as follows. For T given triplets in the form of (Anchor, Positive, Negative), where the first two are (the multi-omics data of) responder cell lines to a given anti-cancer drug and the last one is (the multi-omics data of) a non-responder to that drug, we require the following condition:

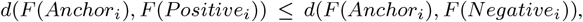
where *d(.)* is an arbitrary distance function—we used the Euclidean distance.

If we move the right hand-side to the left, we obtain:

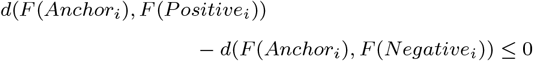

In order to avoid the trivial zero solution, a margin *ξ* > 0 is required:

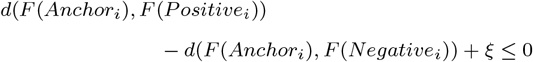

We want the distance of the Anchor and the Negative to be larger than the distance of the Anchor and the Positive. Thus, the value of the triplet loss function for the i-th triplet is:

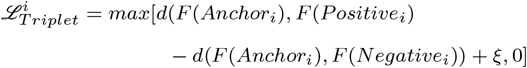

and the total triplet loss for T triplets is:

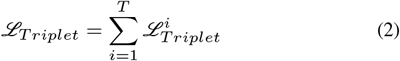

Generally, there are two approaches to select triplets for the triplet loss function: offline selection and online selection. The offline selection builds the triplets based on the value of the labels (in this case the drug response) before training the model. The online selection selects the triplets from samples in each mini-batch during the training. We adopted the online approach. Triplets can be built based on all possible combinations of the input samples/mini-batches (soft selection) or based only on those triplets with high triplet loss value (Hard selection). Soft selection provides the model with more training triplet examples but the network might rely too much on easy cases, and as a result may be unable to perform well on hard examples (Schroff *et al.*, 2015). Hard selection solves this problem by only relying on the hard cases in the train data to build the triplets, but this approach may suffer from having fewer training triplets especially in the case of small unbalanced datasets. We adopted the soft selection approach.

Therefore, the combined cost *J* is defined as follows:

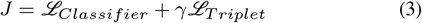
where *γ* is a regularization term for the triplet loss.

### 2.2 Transfer learning for targeted drugs

For targeted drugs, we use transfer learning and train MOLI with a new pan-drug input. This pan-drug input consists of multi-omics profiles and drug responses for a family of targeted drugs that target the same pathway or molecule. Such drugs are expected to produce highly correlated responses in cell lines. One MOLI model is trained for a family of drugs instead of one separate model for each individual drug. This approach increases the training dataset size, since the set of the screened cell lines and the obtained responses are similar but not identical for the drugs of one family. In our experiments, we evaluate transfer learning for EGFR pathway inhibitors due to the availability of external validation data, but the approach is applicable to any family of targeted drugs. Figure 2-B illustrates the idea of transfer learning for targeted drugs.

### 2.3 Predicting drug response for TCGA patients

To study MOLI’s performance, similar to (Geeleher *et al.*, 2017), we employ the model trained on the pan-drug input for the EGFR inhibitors to predict the drug response for patients in several TCGA datasets for which there was no drug response recorded. Since these drugs target EGFR pathway, we expect the expression status of the genes of this pathway to be strongly correlated with the predicted drug response. We obtain the list of genes in EGFR pathway from REACTOME. To study the correlation, we employ multiple linear regression between the predicted responses and the level of expression. We obtain p-values for each gene and correct them for multiple comparison, using Bonferroni correction (*α* = 0.05).

### 2.4 Datasets

We use four main resources in this paper:

- Genomics of Drug Sensitivity in Cancer (GDSC) cell lines dataset (Iorio *et al.*, 2016)
- Patient-Derived Xenograft (PDX) Encyclopedia dataset (Gao *et al.*, 2015)
- TCGA patients with the drug response available in their records (Ding *et al.*, 2016)
- TCGA patients without the drug response (Weinstein *et al.*, 2013)

The GDSC dataset (Yang *et al.*, 2012; Iorio *et al.*, 2016) has created a multi-omics dataset of more than a thousand cell lines from different cancer types, screened with 265 targeted and chemotherapy drugs. We use GDSC as the training dataset due to a high number of screened drugs. Multi-omics profiles and drug responses for GDSC are retrieved from ftp://ftp.sanger.ac.uk/pub/project/cancerrxgene/releases/release-7.0/.

We use the other publicly available multi-omics datasets for validation as follows:

1. We apply PDX Encyclopedia mice models published by (Gao *et al.*, 2015). This dataset has more than 300 PDX models for different cancer types, screened with 34 targeted and chemotherapy drugs.
2. TCGA (Weinstein *et al.*, 2013) data including profiles of tumor samples collected from more than ten thousand patients with different cancer types, downloaded from Firehose Broad GDAC (https://doi.org/10.7908/C11G0KM9, http://gdac.broadinstitute.org/runs/stddata2016_01_28/). For TCGA datasets, we use clinical annotations of the drug response for some patients which were obtained from supplementary material of (Ding *et al.*, 2016).
3. We also use TCGA patients for breast (BRCA), bladder (BLCA), pancreatic (PAAD), lung (LUAD), kidney (KIRP), and prostate (PRAD) cancers. These patients are without the drug response in their records. We note that we used only those genes which are in common for all of the omics data types in both training and validation datasets for each drug. The data preprocessing steps and the used reference files are summarized below and in table 1, and are presented in more detail in the supplementary material (table S1).

**Table 1.** List of the studied drugs from the used resources with multi-omics profiles available.

| Drug | Type | Resource | Number of samples |
| --- | --- | --- | --- |
| Afatinib | Targeted | GDSC | 828 (NR:678, RS:150) |
| Cetuximab | Targeted | GDSC | 856 (NR:735, RS:121) |
| Cetuximab | Targeted | PDX | 60 (NR:55, RS:5) |
| Cisplatin | Chemotherapy | GDSC | 829 (NR:752, RS:77) |
| Cisplatin | Chemotherapy | TCGA | 66 (NR:6, RS:60) |
| Docetaxel | Chemotherapy | GDSC | 829 (NR:764, RS:65) |
| Docetaxel | Chemotherapy | TCGA | 16 (NR:8, RS:8) |
| Erlotinib | Targeted | GDSC | 362 (NR:298, RS:64) |
| Erlotinib | Targeted | PDX | 21 (NR:18, RS:3) |
| Gefitinib | Targeted | GDSC | 825 (NR:710, RS:115) |
| Gemcitabine | Chemotherapy | PDX | 25 (NR:18, RS:7) |
| Gemcitabine | Chemotherapy | TCGA | 57 (NR:36, RS:21) |
| Lapatinib | Targeted | GDSC | 387 (NR:326, RS:61) |
| Paclitaxel | Chemotherapy | GDSC | 389 (NR:363, RS:26) |
| Paclitaxel | Chemotherapy | PDX | 43 (NR:38, RS:5) |

NR: Non-responder; RS: Responder

#### 2.4.1 Gene expression profiles

Raw intensities are obtained from ArrayExpress (E-MTAB-3610) for GDSC dataset were RMA-normalized (Irizarry *et al.*, 2003), log-transformed and aggregated to the level of genes. Gene expression values of PDX and all TCGA datasets are converted to TPM (Li and Dewey, 2011) and log-transformed. FPKM values for PDX samples are converted into TPM and log-transformed. To make expression profiled by different platforms comparable, we standardize gene expression and perform pairwise homogenization procedure, as described in (Johnson *et al.*, 2007; Geeleher *et al.*, 2014). Also, in each dataset we exclude the 5% of genes with lowest variance assuming them to be not informative.

#### 2.4.2 Somatic copy number profiles

We remove unreliable segments from genome segmentation files for TCGA datasets and assign every gene a value corresponding to the intensity log-ratio of the segment it overlaps. If the gene overlaps more than one segment, we keep the most extreme log-ratio value. Different from TCGA, the GDSC and PDX datasets provided gene-level estimates of total copy number. In order to make these data comparable with TCGA, we compute for every gene the logarithm of its copy number divided by the ploidy of copy-neutral state in the sample. Finally, for all four datasets we binarize gene-level copy number estimates assigning zeros to copy-neutral genes and ones to all genes overlapping deletions or amplifications. .

#### 2.4.3 Somatic point mutations

Similarly with previous works (Iorio *et al.*, 2016; Ding *et al.*, 2018), we assign ones to genes carrying somatic point mutations and zeros to all others.

## 3 Results

### 3.1 Experimental Design

In our experiments, we investigated the following questions:

- Does MOLI outperform single-omics and early integration baselines in terms of prediction AUC on PDX and patient data?
- Does transfer learning work for targeted drugs, i.e. does MOLI trained on pan-drug data outperform MOLI trained on drug-specific (single drug) data?
- Finally, for the targeted drugs, does the predicted response by MOLI have associations with the target of that drug?

We trained MOLI on GDSC cell lines screened with Docetaxel, Cisplatin, Gemcitabine, Paclitaxel, Erlotinib, and Cetuximab. We chose these drugs based on availability of PDX/patient multi-omics data for these drugs which is necessary for external validations. We trained all of the baselines for the same drugs and compared them to MOLI in terms of prediction AUC.

We compared MOLI against early integration via deep neural networks inspired by (Ding *et al.*, 2018), against the single-omics (gene expression) ridge regression method proposed by (Geeleher *et al.*, 2014), against an ordinary feed forward network with classification loss trained on the expression data, and against a version of MOLI trained only on the gene expression data. To test whether the triplet loss contributes to improve the performance, we compared MOLI to a late integration feed forward network with an architecture similar to MOLI but using only a classification loss.

Finally, to study transfer learning for the targeted drugs, we focused on drugs that target the EGFR pathway because we have Cetuximab and Erlotinib that target this pathway in the PDX dataset utilized for external validations. In addition, GDSC was screened with numerous drugs that target EGFR including: Afatinib, Cetuximab, Erlotinib, Gefitinib, and Lapatinib. We used multi-omics data of all of these drugs in GDSC and created a large training set (>3,000 samples). We trained MOLI on this pan-drug data and compared the results to MOLI which was trained on the drug-specific inputs.

We used 5-fold cross validation in most of the experiments to tune the hyper-parameters of the deep neural networks based on the AUC. The hyper-parameters tuned were number of nodes in the hidden layers, learning rates, mini-batch size, weight decay, the dropout rate, number of epochs, and margin and regularization term (only for the triplet loss).

Details on the ranges considered for each hyper-parameter and the selected settings for each drug are provided in tables S2 and S3 in the supplementary material. Finally, the network was re-trained with the obtained hyper-parameters on the entire dataset for that drug (train and validation). We used Adagrad for optimizing parameters in all of the deep neural networks (Duchi *et al.*, 2011). We used the Pytorch framework to implement all deep neural networks codes. For the ridge regression pipeline, we downloaded the implemented pipeline with leave-one-out cross validation provided by the original authors (Geeleher *et al.*, 2014) and applied it to our datasets. To make sure that both the downloaded pipeline and the way we preprocessed the gene expression data are correct, we evaluated it on the datasets from the original paper and got AUCs for Docetaxel and Bortezomib comparable to those of (He *et al.*, 2018).

### 3.2 Multi-omics integration by MOLI improves the drug response performance

Table 2 reports the performance of MOLI and the baselines in terms of AUC. The complete MOLI, i.e. MOLI trained on multi-omics data and using its combined cost function achieved the best performance in three (out of seven) drugs (Paclitaxel, Cetuximab, and Erlotinib) and the second best performance for three drugs (PDX Gemcitabine, TCGA Gemcitabine, and Cisplatin). The complete MOLI also achieved the best performance for Cisplatin when it was trained only on the gene expression data. Multi-omics early integration had the best performance for Gemcitabine in PDX data, the ordinary feed forward network achieved the best performance for Docetaxel, and MOLI with only the classification loss had the best performance for Gemcitabine in TCGA data. MOLI when trained on the pan-drug input (only applicable for targeted drugs), had significantly better performance compared to itself when it was trained on the drug-specific inputs for Erlotinib and Cetuximab. The majority of the baselines had either poor performance or no stable classifier status (NSC) for Paclitaxel and Erlotinib. NSC means that during either cross validation or final re-training with the obtained hyper-parameters the cost and/or AUC curves were fluctuating. This may be due to the small number of samples, because both of these drugs had the fewest number of cell lines (*∼*400). Also, we observed NSC in the early integration baseline for four drugs which may be due to the concatenation at the beginning because it increased the dimensionality substantially, which makes feature learning harder for the autoencoder and later the classifier in this method.

**Table 2.** Performance of different versions of MOLI compared to the baselines in terms of prediction AUC across two targeted therapeutics and five chemotherapy agents.

| Method/<br>Drug | PDX<br>Paclitaxel | PDX<br>Gemcitabine | PDX<br>Cetuximab | PDX<br>Erlotinib | TCGA<br>Docetaxel | TCGA<br>Cisplatin | TCGA<br>Gemcitabine | Input<br>omics |
| --- | --- | --- | --- | --- | --- | --- | --- | --- |
| (Geeleher *et al.*, 2014) | 0.52 | 0.59 | <u>0.58</u> | <u>0.67</u> | 0.59 | 0.62 | 0.53 | Expression |
| Early integration | NSC | **0.66** | NSC | NSC | 0.52 | NSC | 0.59 | Multi |
| Feed Forward Net | 0.68 | 0.48 | 0.43 | 0.37 | **0.69** | 0.44 | <u>0.65</u> | Expression |
| MOLI complete | <u>0.69</u> | 0.52 | 0.51 | 0.39 | <u>0.63</u> | **0.75** | 0.64 | Expression |
| MOLI with classifier | NSC | 0.55 | 0.46 | NSC | 0.58 | 0.6 | **0.69** | Multi |
| MOLI complete | **0.74** | <u>0.64</u> | 0.53 | 0.63 | 0.58 | <u>0.66</u> | <u>0.65</u> | Multi |
| MOLI complete pan-drug | NA | NA | **0.80** | **0.72** | NA | NA | NA | Multi |

NSC: no stable classifier during cross validation or final training. **boldface** indicates the best method for the corresponding drug and underline indicates the second best method. NA: the corresponding drug is not for targeted therapy. Complete: when MOLI has both classification and the triplet losses.

MOLI achieved an AUC of greater than 0.7 for four drugs (Paclitaxel, Cetuximab, Erlotinib, and Cisplatin) which may be beneficial for precision oncology particularly for the targeted drugs (Cetuximab and Erlotinib).

### 3.3 Transfer learning for targeted drugs improves performance significantly

We observed that for the targeted drugs (in our experiment, EGFR inhibitors), MOLI trained on the pan-drug multi-omics inputs achieved significantly better performance than MOLI trained on drug-specific inputs. Pan-drug MOLI achieved an AUC of 0.8 for Cetuximab and 0.72 for Erlotinib which were significantly higher than the drug-specific performance. This suggests that transfer learning can improve the prediction performance for the targeted drugs.

### 3.4 Predictions for TCGA patients by MOLI have associations with EGFR genes

We applied MOLI (trained on the pan-drug input for EGFR inhibitors) to multi-omics data without drug response downloaded from TCGA (breast, bladder, pancreatic, lung, kidney, and prostate cancers) and predicted the response for these patients. According to the p-values obtained from multiple linear regression, there are a number of strong associations between EGFR genes and the responses predicted by MOLI. For breast cancer, we observed statistically significant associations between the level of expression in AP2A1 (*P* = 0.007), CALM2 (*P* = 0.01), CLTA (*P* = 0.0002), EGFR (*P* = 1 × 10^−5^), PIK3CA (*P* = 0.007), and UBA52 (*P* = 3 × 10^−6^) genes and the predicted responses. For prostate cancer, we found that the predicted responses have statistically significant associations with the expression of AKT1 (*P* = 0.02), CDK1 (*P* = 0.01), RICTOR (*P* = 0.0002), CREB1 (*P* = 0.02), and CSK (*P* = 0.01). In kidney cancer, expression of EGFR (*P* = 0.04) gene had association with the predicted response. In lung cancer, we observed significant associations for CDC42 (*P* = 0.04), EGFR (*P* = 3 × 10^−5^), and PRKAR2A (*P* = 0.01) genes. However, for bladder and pancreatic cancers, we did not observe any significant associations.

## 4 Discussion

In this paper, we proposed MOLI, a Multi-Omics Late Integration method based on deep neural networks to predict drug response. MOLI integrates somatic mutation, CNA, and gene expression data and predicts the drug responses. To the best of our knowledge, MOLI is the first end-to-end method for multi-omics late integration with deep neural networks that utilizes a combined cost function. Our experiments showed that MOLI with its combined cost function can achieve better performance than single-omics and early integration multi-omics methods based on deep neural networks. We also observed that transfer learning for targeted drugs improves the prediction performance compared to drug-specific inputs. To the best of our knowledge, this is the first method to use transfer learning with a pan-drug approach for targeted drugs. Finally, we analyzed MOLI’s predictions for drugs targeting the EGFR pathway on breast, kidney, lung, and prostate cancer patients in TCGA. We showed that MOLI’s predictions have statistically significant associations with the level of expression for some of the genes in the EGFR pathway, including the EGFR gene itself, for breast, kidney, and lung cancers.

We would like to point out the following directions for future research: Although we used only somatic mutation, CNA, and gene expression data in our experiments, MOLI can be extended for integrating other omics data types. For example, proteomics data can be a good candidate because it has been shown to be a contributing factor in pan-cancer drug response prediction (Ali *et al.*, 2017) and is known to be in concordance with the other omics data types (Ryan *et al.*, 2017; Gonçalves *et al.*, 2017). We performed experiments on transfer learning only for the drugs that target EGFR, but this approach is also applicable for other families of targeted drugs if multi-omics data are available for external validation. Another advantage of the pan-drug approach is that there is no need to train separate pan-drug models for each EGFR inhibitor, and one model can be validated on different external datasets. In the drug-specific approach, we trained one model on Cetuximab data and another one on Erlotinib data, and could not validate them on each other’s external validation data. However, in the pan-drug approach, we trained one model for all of the EGFR inhibitors and validated it on both Cetuximab and Erlotinib data.

While we studied only the triplet loss for optimizing the concatenated representation, we note that this loss function can be replaced by other similar losses such as the contrastive loss function which was used in the Siamese network (Hadsell *et al.*, 2006). We trained separate MOLI models for different drugs, but it is an interesting direction for future research to utilize multi-task learning (Yuan *et al.*, 2016) and predict the outcome for multiple drugs at the same time.

In all of the experiments and utilized datasets, we used pan-cancer inputs. The advantage of using pan-cancer multi-omics input is that it can address, to some degree, the challenge of intertumor heterogeneity (Almendro *et al.*, 2013). However, these datasets are not suitable for addressing intratumor heterogeneity, which would require other resources such as single cell data.

We would like to point out the following limitations of this study:

1. The datasets used were from different resources were not in the same format and required substantial preprocessing and standardization (see the supplementary material). For example, different studies used different pipelines to detect CNA and reported different estimates of copy number which could not be compared directly. A similar issue was also observed for the drug response. While the GDSC cell lines used IC50 as the response measure, the majority of datasets used other metrics to measure the response. For example, the PDX dataset used tumor volume based on RECIST criteria to define responders and non-responders. Therefore, lack of standardization on both the input and the output side adds extra challenges to the drug response prediction task.
2. In this study we focused on monotherapy and did not explore the effect of the combination of drugs.
3. We did not discriminate between driver and passenger events in the somatic mutation and CNA data and treated all of them similarly. However, in reality, the majority of these genomic alterations seem to have no impact on cancer development (Vogelstein *et al.*, 2013) and might appear just by chance. Therefore, in future work, we plan to use another format for these data types to distinguish between potential driver and passenger events.
4. All of the datasets used suffered from severely unbalanced class distributions, since the number of responders was much smaller than the number of non-responders. We addressed this problem by oversampling the minority class. However, this approach often causes overfitting particularly for deep neural networks with many parameters. We reduced overfitting with strong regularization such as high dropout rate and weight decay. Moreover, using triplets as input of the network increased the number of samples and led to a more stable network, due to the large number of different combinations for triplets.

## 5 Conclusion

In this paper, we proposed MOLI, a method for drug response prediction based on deep neural networks and Multi-Omics Late Integration. We trained MOLI on a pan-cancer cell line dataset and successfully validated it on PDX and patient data for five chemotherapy agents and two targeted therapeutics.

Our results suggest four major findings:

1. MOLI outperforms single-omics (gene expression) prediction performance in terms of AUC.
2. MOLI outperforms deep neural networks using early integration in terms of AUC.
3. MOLI with its combined cost function outperforms single- and multi-omics baselines with only the classification loss.
4. MOLI trained on the pan-drug inputs, employing transfer learning, outperforms MOLI trained on drug-specific inputs for targeted therapeutics that target EGFR.

Finally, we analyzed the biological significance of MOLI and found substantial evidence that the responses predicted by MOLI have statistically significant associations with the expression level of numerous genes in the EGFR pathway for TCGA patients with breast, kidney, lung, and prostate cancers.

In conclusion, our experimental results suggest MOLI may have a role in precision oncology where currently only *∼*5% of all patients benefit from precision oncology.

## Acknowledgements

We would like to thank Hossein Asghari, Baraa Orabi, Raunak Shrestha, and Yen-yi Lin (Vancouver Prostate Centre), Anne-Marie Therien-Daniel, Nazanin Mehrasa, and Mehrdad Mansouri (Simon Fraser University), Paul Geeleher (St. Jude Children’s Research Hospital), and Lukas Folkman (CeMM) for their kind supports. We also would like to thank Compute Canada, WestGrid, and Vancouver Prostate Centre for providing us with the computational resources for this research.

## Authors’ contributions

Study concept and design: H.SN., C.C.C., M.E.

Deep learning design, implementations, and analysis: H.SN.

Data preprocessing, analysis, and interpretation: O.Z.

Analysis and interpretation of results: H.SN., O.Z.

Drafting of the manuscript: All authors read and approved the final manuscript.

Supervision: C.C.C., M.E.

