## Supplementary material for "MOLI: Multi-Omics Late Integration with deep neural networks for drug response prediction": Supp Material

**Hossein Sharifi-Noghabi[1,3], Olga Zolotareva[2], Colin C. Collins[3,4], and Martin Ester[1,3]**
[1]School of Computing Science, Simon Fraser University, Burnaby, BC, Canada
[2]International Research Training Group "Computational Methods for the Analysis of the Diversity and Dynamics of Genomes" and Genome Informatics, Faculty of Technology and Center for Biotechnology, Bielefeld University, Germany
[3]Vancouver Prostate Centre, Vancouver, BC, Canada
[4]Department of Urologic Sciences, University of British Columbia, Vancouver, BC, Canada

#### Preprocessing steps

##### Gene expression profiles

Raw CEL files for GDSC cohort were obtained from ArrayExpress website (https://www.ebi.ac.uk/arrayexpress/E-MTAB-3610). RMA (robust multi-array average) normalization (Irizarry *et al.*, 2003) of raw intensities was done using *justRMA()* function from affy (v1.54.0) R package. This function performs background correction, quantile normalization, and log-transformation of probe intensities. CDF library files and probe set annotations for corresponding array platforms were obtained from BrainArray (Dai *et al.*, 2005) v22.0.0 (http://brainarray.mbni.med.umich.edu). After the normalization, probe set identifiers were mapped to Entrez Gene identifiers. Intensities of the probe set corresponding to a single gene were summarized using *collapseRows()* function (Miller *et al.*, 2011) from WGCNA (v 1.64.1) R package with method="Average". Probe sets mapped to more than one Entrez gene were considered unspecific and removed.

For all TCGA cohorts, we used the estimated fractions of transcripts computed by RSEM method (Li and Dewey, 2011) (scaled_estimates) provided by Firehose Broad GDAC (http://gdac.broadinstitute.org/runs/stddata__2016_01_28/data/), multiplied by $10^6$ to obtain TPM (Li and Dewey, 2011) and log2-transformed. FPKM values for PDX samples were obtained from the supplementary table published by (Gao *et al.*, 2015), converted into TPM, and log-transformed log2(TPM+1).

$$TPM_i = \frac{FPKM_i}{\sum_j FPKM_j} * 10^6$$ (Pachter, 2011)

Gene symbols were mapped to current Entrez Gene IDs using the table provided by NCBI (tp.ncbi.nih.gov:gene/DATA/GENE_INFO/Mammalia/Homo_sapiens.gene_info.gz).

To make expression measures in different datasets comparable, we standardized gene expressions within each cohort and performed pairwise homogenization procedure, as described in (Johnson *et al.*, 2007; Geeleher *et al.*, 2014). Briefly, for every pair of training and testing datasets, we kept only genes presenting in both datasets and applied *ComBat()* function (Johnson *et al.*, 2007) from SVA R package v3.24.4. Finally, for each dataset, we excluded 5% of genes with the lowest variance assuming them not informative.

**Copy number profiles**

In all TCGA cohorts, copy numbers were profiled by Affymetrix SNP6.0 arrays. Probe intensities measured for a sample were normalized by intensities in the most similar normal samples from HapMap (Johnson *et al.*, 2007; Geeleher *et al.*, 2014; International HapMap 3 Consortium *et al.*, 2010) and log2-transformed. The resulted point estimates of intensity log-ratios (logR) were united into segments with the same level of logR using the circular binary segmentation (CBS) algorithm (Olshen *et al.*, 2004). The resulted genome segmentation files for TCGA cohorts were downloaded from Firehose Broad GDAC (data published on 2016_01_28). These files contained hg19 coordinates of segments, a number of probes united into a segment, and an averaged intensity log-ratios reflecting the ratio of DNA amount in these segment to the DNA amount in the copy-neutral state. Although for TCGA we used segmentation files with "masked" putative germline CNAs detected in a panel of normals, we noticed that many tumor samples still contained some segments matching with segments in normals derived from the same patient. This might be either due to a cross-sample contamination when the normal sample was mixed with tumor DNA, or the result of the inclusion of sample-specific germline CNA into somatic CNA profile of the tumor. To remove likely germline segments from tumor CNA profiles, we performed two additional steps of filtering for TCGA samples. First, we excluded all segments with logR below 0.46 and above -0.68 from matched normal CNA profiles. These thresholds corresponded to one copy gain and loss and -1 copy in 75% of a normal cell. We selected these thresholds based on the assumption that if tumor content in a matched normal sample is not high by applying these thresholds we exclude putative tumor CNAs from normal samples. Second, we compared the remaining segments in normal profiles with tumor profiles and removed all tumor segments covered by more than 80% by normal segments. Segments including less than five probes removed from all CNA profiles, assuming that such segments are noisy. Finally, we overlapped remained segments with gene annotation for GRCh37/hg19 assembly obtained from NCBI and assigned every gene a value corresponding to logR of the segment it overlaps. If the gene overlapped more than one segment, we kept the most extreme log-ratio value. Genes overlapped no segments or only segments with logR below 0.20 or above -0.23 were considered to be copy-neutral. These thresholds correspond to log-ratios of 1-copy gain and 1-copy loss respectively occurred in 30% of cells.

GDSC and PDX datasets were obtained from ftp://ftp.sanger.ac.uk/pub/project/cancerrxgene/releases/release-7.0/Gene_level_CN.xlsx and supplementary files from (Gao et al. 2015), respectively. In contrast with TCGA, these projects provided gene-level estimated total copy numbers (CN). In order to make these data comparable with TCGA, we computed for every gene the logarithm of its CN divided by ploidy of copy-neutral state in the sample. Copy-neutral state was predicted for each sample based on the distribution of gene-level CN estimates, assuming that the mode closest to the median corresponds to the copy-neutral state. Similarly, with TCGA, all genes with log-ratios below 0.2 or above -0.23 were assumed to be neutral. Finally, for all four cohorts, we binarized gene-level CN estimates assigning zero to copy-neutral genes and one to all genes overlapping deletions or amplification.

**Point mutations**

Somatic point mutations in GDSC cell lines were retrieved from ftp://ftp.sanger.ac.uk/pub/project/cancerrxgene/releases/release-7.0/WES_variants.xlsx. MAF files for TCGA samples from all cohorts were downloaded from http://gdac.broadinstitute.org/runs/stddata__2016_01_28/data/. List of somatic mutations in PDX samples was obtained from supplementary tables (Gao *et al.*, 2015), tab "pdxe_mut_and_cn2". Amplification and deletions were removed. From all reported point mutations, we selected only those affecting protein structure and filtered out silent ones. Similarly, with previous works (Iorio *et al.*, 2016);(Geeleher *et al.*, 2014; Ding *et al.*, 2018), we assigned one to genes carrying any nonsynonymous somatic mutations and zero to all others. All gene IDs were mapped to Entrez Gene IDs.

### Supplementary tables

| Table S1 Drug responses available for GDSC, TCGA and PDX cohorts. | | | |
|---|---|---|---|
| **cohort** | **sources** | **original response measure** | **response interpretation** |
| **GDSC (binary response)** | Binary response: TableS5C.xlsx from (Iorio *et al.*, 2016) | RS –Non-responder, RS – Responder; | - |
| **GDSC (continuous response)** | log(IC50): TableS4A.xlsx from (Iorio *et al.*, 2016) | log(IC50) | - |
| **PDX** | (Gao *et al.*, 2015) Supplementary file nm.3954-S2.xlsx, tab “PCT curve metrics”, ResponseCategory field | RECIST Response Categories | CR and PR are considered as sensitive and SD and PD of the entries are considered as resistant; Unstable responses were excluded as well as response to combo treatment |
| **TCGA** | (Ding *et al.*, 2016), Supplementary Table S2 | RECIST Response Categories | CR and PR are considered as sensitive and SD and PD of the entries are considered as resistant; Only single drug treatments kept |

| Table S2 Considered ranges for each hyper-parameter for cross validation | |
|---|---|
| **Hyper_parameter** | **Range** |
| Mini-batch size | [8, 16, 32, 64]* |
| Number of nodes | [2048, 1024, 512, 256, 128, 64, 32, 16] |
| Margin | [0.5, 1, 1.5, 2, 2.5, 3, 3.5] |
| Learning rate | [0.1, 0.5, 0.01, 0.05, 0.001, 0.005, 0.0001, 0.0005, 0.00001, 0.00005] |
| Number of epochs | [5, 10, 20, 30 ,40, 50, 60, 70, 80, 90, 100, 150, 200] |
| Dropout rate | [0.3, 0.4, 0.5, 0.6, 0.7, 0.8] |
| Weight decay | [0.1, 0.01, 0.001, 0.1, 0.0001] |
| Gamma | [0.1, 0.2, 0.3, 0.4, 0.5, 0.6] |
| * In order to make sure each mini-batch has at least three members to form the triplets,<br>for some of the drugs we had to change the size to 13, 14, 30, 36, 60, and 62. | |

**Table S3 Obtained hyper-parameters based on cross validation**

| Methods for Paclitaxel | mini-batch size | #nodes | learning rate expression | learning rate mutation | learning rate CNA | Learning rate Classifier | dropout expression | dropout mutation | dropout CNA | weight decay | dropout classifier | gamma | #epoch | #Folds | margin |
|---|---|---|---|---|---|---|---|---|---|---|---|---|---|---|---|
| AE Early integration | NSC | NSC | NSC | NSC | NSC | NSC | NSC | NSC | NSC | NSC | NSC | NSC | NSC | 7,10 | NSC |
| Feed Forward | 13 | 128 | 0.001 | NA | NA | 0.05 | 0.5 | NA | NA | 0.01 | 0.3 | NA | 10 | 5 | NA |
| MOLI_Complete_OnlyExprs | 36 | 64 | 0.05 | NA | NA | 0.005 | 0.5 | 0.5 | 0.5 | 0.001 | 0.3 | 0.005 | 10 | 5 | 1.5 |
| MOLI_OnlyClassificationLoss | NSC | NSC | NSC | NSC | NSC | NSC | NSC | NSC | NSC | NSC | NSC | NSC | NSC | 7 | NSC |
| MOLI_Complete | 64 | 512-256-1024* | 0.0005 | 0.5 | 0.5 | 0.5 | 0.4 | 0.4 | 0.5 | 0.0001 | 0.3 | 0.6 | 10 | 5 | 0.5 |
| **Methods for PDX Gemcitabine** | **mini-batch size** | **#nodes** | **learning rate expression** | **learning rate mutation** | **learning rate CNA** | **Learning rate Classifier** | **dropout expression** | **dropout mutation** | **dropout CNA** | **weight decay** | **dropout classifier** | **gamma** | **#epoch** | **#Folds** | **margin** |
| Early integration | 62 | 256,128 | NA | NA | NA | 0.05 | NA | NA | NA | 0.001 | 0.2 | NA | 10 | 7 | NA |
| Feed Forward | 30 | 1024 | 0.05 | NA | NA | 0.001 | 0.5 | NA | NA | 0.1 | 0.3 | NA | 10 | 5 | NA |
| MOLI_Complete_OnlyExprs | 64 | 32 | 0.1 | NA | NA | 1.00E-05 | 0.5 | NA | NA | 0.1 | 0.3 | 0.1 | 10 | 5 | 2.5 |
| MOLI_OnlyClassificationLoss | 62 | 1024,64** | 0.1 | 5.00E-05 | 0.01 | 0.005 | 0.5 | 0.5 | 0.5 | 0.01 | 0.4 | NA | 5 | 5 | NA |
| MOLI_Complete | 13 | 256,32,64 | 0.05 | 1.00E-05 | 0.0005 | 0.001 | 0.4 | 0.6 | 0.3 | 0.01 | 0.6 | 0.3 | 5 | 5 | 1.5 |
| **Methods for Cetuximab** | **mini-batch size** | **#nodes** | **learning rate expression** | **learning rate mutation** | **learning rate CNA** | **Learning rate Classifier** | **dropout expression** | **dropout mutation** | **dropout CNA** | **weight decay** | **dropout classifier** | **gamma** | **#epoch** | **#Folds** | **margin** |
| Early integration | NSC | NSC | NSC | NSC | NSC | NSC | NSC | NSC | NSC | NSC | NSC | NSC | NSC | 7,10 | NSC |
| Feed Forward | 30 | 128 | 0.05 | NA | NA | 0.5 | 0.5 | NA | NA | 0.1 | 0.3 | NA | 10 | 5 | NA |
| MOLI_Complete_OnlyExprs | 16 | 512 | 0.001 | NA | NA | 5.00E-05 | 0.5 | 0.5 | 0.5 | 0.001 | 0.5 | 0.1 | 10 | 5 | 2 |
| MOLI_OnlyClassificationLoss | 32 | 1024-128 | 1.00E-05 | 0.0005 | 0.0001 | 5.00E-05 | 0.5 | 0.5 | 0.5 | 0.001 | 0.4 | NA | 10 | 7 | NA |
| MOLI_Complete | 30 | 256,512,128 | 0.0001 | 0.0005 | 0.0005 | 0.0005 | 0.3 | 0.8 | 0.8 | 0.01 | 0.4 | 0.2 | 10 | 5 | 2 |
| MOLI_Complete_Pan_Drug | 16 | 32,16,256* | 0.001 | 0.0001 | 5.00E-05 | 0.005 | 0.5 | 0.8 | 0.5 | 0.0001 | 0.3 | 0.5 | 20 | 5 | 1.5 |
| **Methods for Erlotinib** | **mini-batch size** | **#nodes** | **learning rate expression** | **learning rate mutation** | **learning rate CNA** | **Learning rate Classifier** | **dropout expression** | **dropout mutation** | **dropout CNA** | **weight decay** | **dropout classifier** | **gamma** | **#epoch** | **#Folds** | **margin** |
| Early integration | NSC | NSC | NSC | NSC | NSC | NSC | NSC | NSC | NSC | NSC | NSC | NSC | NSC | 7,10 | NSC |
| Feed Forward | 14 | 512 | 0.0001 | NA | NA | 0.001 | 0.5 | NA | NA | 0.0001 | 0.4 | NA | 10 | 5 | NA |
| MOLI_Complete_OnlyExprs | 64 | 1024 | 0.001 | NA | NA | 0.1 | 0.5 | NA | NA | 0.0001 | 0.5 | 0.5 | 10 | 5 | 1 |
| MOLI_OnlyClassificationLoss | NSC | NSC | NSC | NSC | NSC | NSC | NSC | NSC | NSC | NSC | NSC | NSC | NSC | 5,7,10 | NSC |
| MOLI_Complete | 32 | 64 | 0.5 | 0.5 | 0.1 | 0.1 | 0.5 | 0.5 | 0.5 | 0.01 | 0.5 | 0.6 | 5 | 5 | 1 |
| MOLI_Complete_Pan_Drug | 16 | 32,16,256* | 0.001 | 0.0001 | 5.00E-05 | 0.005 | 0.5 | 0.8 | 0.5 | 0.0001 | 0.3 | 0.5 | 20 | 5 | 1.5 |
| **Methods for Docetaxel** | **mini-batch size** | **#nodes** | **learning rate expression** | **learning rate mutation** | **learning rate CNA** | **Learning rate Classifier** | **dropout expression** | **dropout mutation** | **dropout CNA** | **weight decay** | **dropout classifier** | **gamma** | **#epoch** | **#Folds** | **margin** |
| Early integration | 60 | 256,128 | NA | NA | NA | 0.005 | NA | NA | NA | 0.001 | 0.2 | NA | 15 | 5 | NA |
| Feed Forward | 64 | 128 | 1.00E-04 | NA | NA | 5.00E-05 | 0.5 | NA | NA | 0.1 | 0.3 | NA | 10 | 5 | NA |
| MOLI_Complete_OnlyExprs | 36 | 32 | 0.1 | NA | NA | 1.00E-05 | 0.5 | NA | NA | 0.0001 | 0.5 | 0.5 | 10 | 5 | 3 |
| MOLI_OnlyClassificationLoss | 60 | 512128** | 0.0001 | 0.001 | 0.01 | 0.005 | 0.5 | 0.5 | 0.5 | 0.001 | 0.5 | NA | 30 | 5 | NA |
| MOLI_Complete | 8 | 16 | 0.0001 | 0.0005 | 0.0005 | 0.001 | 0.5 | 0.5 | 0.5 | 0.001 | 0.5 | 0.4 | 10 | 5 | 0.5 |
| **Methods for Cisplatin** | **mini-batch size** | **#nodes** | **learning rate expression** | **learning rate mutation** | **learning rate CNA** | **Learning rate Classifier** | **dropout expression** | **dropout mutation** | **dropout CNA** | **weight decay** | **dropout classifier** | **gamma** | **#epoch** | **#Folds** | **margin** |
| Early integration | 15 | 2048-128 | NA | NA | NA | 0.01 | NA | NA | NA | 0.01 | 0.2 | NA | 25 | 5 | NA |
| Feed Forward | 64 | 64 | 0.0001 | NA | NA | 0.0001 | 0.5 | NA | NA | 0.001 | 0.5 | NA | 10 | 5 | NA |
| MOLI_Complete_OnlyExprs | 64 | 256 | 0.1 | NA | NA | 0.005 | 0.5 | NA | NA | 0.0001 | 0.5 | 0.5 | 20 | 5 | 3 |
| MOLI_OnlyClassificationLoss | 60 | 256 | 5.00E-05 | 0.0005 | 0.05 | 0.005 | 0.5 | 0.5 | 0.5 | 0.01 | 0.6 | NA | 60 | 5 | NA |
| MOLI_Complete | 15 | 128 | 0.05 | 0.005 | 0.005 | 0.0005 | 0.5 | 0.6 | 0.8 | 0.1 | 0.6 | 0.2 | 20 | 5 | 0.5 |
| **Methods for TCGA Gemcitabine** | **mini-batch size** | **#nodes** | **learning rate expression** | **learning rate mutation** | **learning rate CNA** | **Learning rate Classifier** | **dropout expression** | **dropout mutation** | **dropout CNA** | **weight decay** | **dropout classifier** | **gamma** | **#epoch** | **#Folds** | **margin** |
| Early integration | 32 | 2048-256 | NA | NA | NA | 0.01 | NA | NA | NA | 0.01 | 0.2 | NA | 10 | 5 | NA |
| Feed Forward | 64 | 1024 | 1.00E-05 | NA | NA | 0.0001 | 0.5 | NA | NA | 0.001 | 0.3 | NA | 10 | 5 | NA |
| MOLI_Complete_OnlyExprs | 64 | 1024 | 1.00E-05 | NA | NA | 1.00E-05 | 0.5 | NA | NA | 0.1 | 0.4 | 0.005 | 10 | 5 | 2 |
| MOLI_OnlyClassificationLoss | 62 | 256,16** | 0.1 | 0.1 | 0.05 | 0.005 | 0.5 | 0.5 | 0.5 | 0.1 | 0.3 | NA | 50 | 5 | NA |
| MOLI_Complete | 13 | 16 | 0.001 | 0.0001 | 0.01 | 0.05 | 0.5 | 0.5 | 0.5 | 0.001 | 0.5 | 0.6 | 10 | 5 | 2 |

* #nodes for expression, mutation, and CNA nodes were different

** the classifier has a second hidden layer and the second number is #nodes in that layer

| AutoEncoder for Early integration | mini-batch size | #nodes | learning rate | dropout | #epoch | #Folds |
|---|---|---|---|---|---|---|
| Paclitaxel | 64 | 1024,64 | 0.05 | 0.5 | 40 | 5 |
| Cetuximab | 64 | 1024,64 | 0.1 | 0.5 | 150 | 5 |
| PDX-Gemcitabine | 64 | 256,128 | 0.05 | 0.5 | 100 | 5 |
| Erlotinib | 64 | 2048-128 | 0.005 | 0.5 | 100 | 5 |
| TCGA-Gemcitabine | 64 | 2048-256 | 0.01 | 0.5 | 20 | 5 |
| Cisplatin | 32 | 2048-128 | 0.05 | 0.5 | 200 | 5 |
| Docetaxel | 64 | 256,128 | 0.1 | 0.5 | 20 | 5 |
